# FLOW: maximum flow framework for the identification of factors mediating the signaling convergence of multiple receptors

**DOI:** 10.1101/2022.06.27.497758

**Authors:** Ron Sheinin, Shai Dulberg, Ayelet Kaminitz, Roded Sharan, Asaf Madi

**Affiliations:** Blavatnik School of Computer Science, Tel Aviv University, 69978 Tel Aviv, Israel; Department of Pathology, Sackler Faculty of Medicine, Tel Aviv University, Tel Aviv, Israel

**Author notes:** Equal contribution (Roded Sharan, Asaf Madi).

## Abstract

**Motivation:** Cell-cell crosstalk involves simultaneous interactions of multiple receptors and ligands, followed by downstream signaling cascades working through receptors converging at dominant transcription factors which then integrate and propagate multiple signals into a cellular response. Single cell RNAseq of multiple cell subsets isolated from a defined microenvironment provides us with a unique opportunity to learn about such interactions reflected in their gene expression levels.

**Results:** We developed the FLOW framework to map the potential ligand-receptor interactions between different cell subsets based on a maximum flow computation in a network of protein-protein interactions (PPIs). The maximum flow approach further allows characterization of the intracellular downstream signal transduction from differentially expressed receptors towards dominant transcription factors, therefore, enabling the association between a set of receptors and their downstream activated pathways. Importantly, we were able to identify key transcription factors toward which the convergence of multiple receptor signaling occurs. These identified factors have a unique role in the integration and propagation of signaling following specific cell-cell interactions.

## Introduction

Cellular microenvironments are a complicated heterogeneous assembly of cells wherein their crosstalk often drives cellular response. Single cell RNAseq technology provides us with a snapshot of this transcriptional heterogeneity.

Cell-to-cell interactions require the activation of a receptor by a physical connection of an appropriate ligand. However, it is not trivial to identify such physical interaction using standard transcriptomic data. Thus, current approaches use heuristic methods to try and assess an “interaction potential” between the different cell populations. Indeed, various efforts have been made in recent years to achieve this goal^1^ but the lack of gold standards makes the task of evaluating and comparing the different methods challenging.

Those methods can be roughly divided into those that aim to predict cell-cell interactions based on ligand-receptor expression alone, and those that also seek to detect evidence for intercellular signaling pathways. The latter usually use graph base approaches in order to conclude downstream signaling scores. For example, NicheNet^2^ uses Personalized PageRank (PPR) on the ligand-signaling network in order to calculate the downstream gene activation score. While the former class of methods like CellChat^3^ models the probability of crosstalk between different cell populations via set of ligand-receptor pairs based on gene expression.

Importantly, although cell-cell interactions may involve a single ligand-receptor interaction, in many cases the simultaneous interactions of multiple receptors/ligands are required. In addition, we hypothesize that in a given microenvironment, following cell-cell interaction, downstream signaling progresses via various receptors converge at dominant transcription factors (TFs) that integrate and propagate the multiple signals into a cellular response. To gain insight into potential interactions, we devised an algorithm, FLOW, that maps the expression of ligand/receptor pairs between different cell populations based on single cell transcriptomes. We then use a framework of maximum flow in a protein-protein interaction (PPI) network to characterize the downstream pathways from a set of differentially expressed receptors toward dominant TFs.

We applied our method to a single cell RNAseq dataset from GL261 brain tumor model which contains both infiltrating and resident cells where cell-cell interactions are known to shape the immunosuppressive microenvironment with implications on survival and response to therapies^4^. Our approach was able to highlight likely routes of signal transduction that are upregulated with the receptors, without being limited to a single pathway or an assumption on the size of the downstream pathway. The FLOW framework enables us not only to reveal key ligand-receptor pairs with association to the downstream activation signal transduction but also to pinpoint important transcription factors which aggregate the complex signal going through multiple receptors and pathways and propagate it into a cellular response (**Fig. 1**).

**Figure 1.**
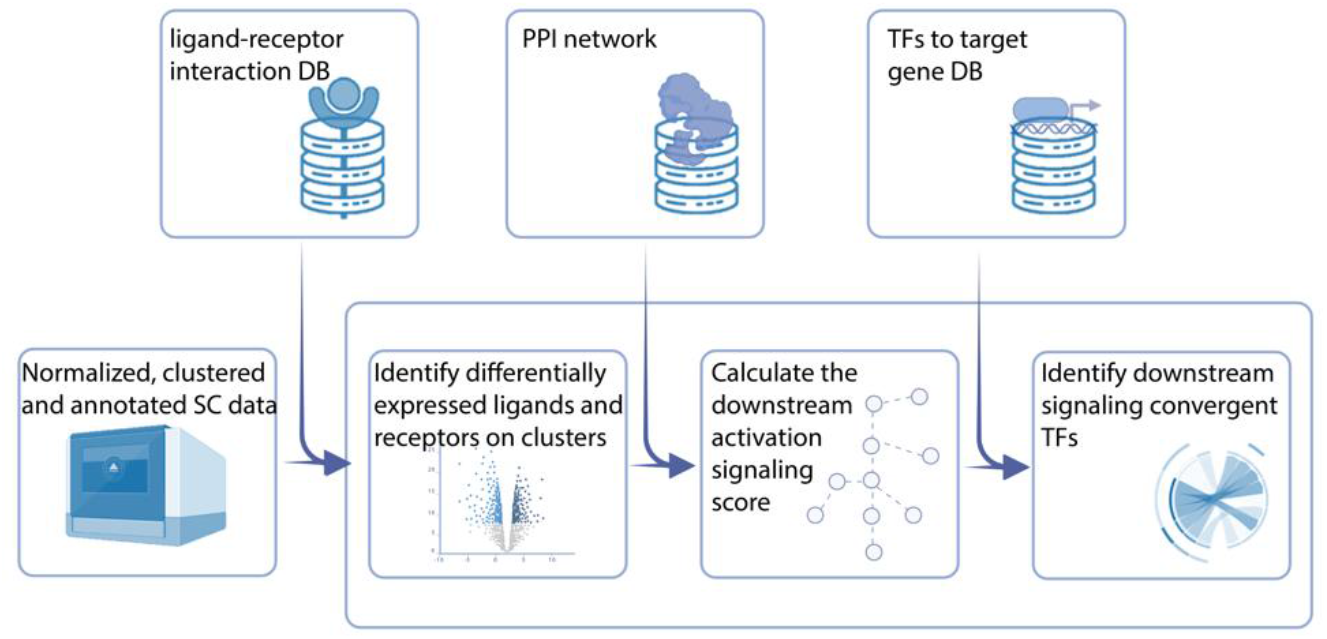
Workflow summary. The FLOW algorithm receives a normalized, clustered and annotated single cell RNAseq dataset. For each pair of cell clusters, a signal sender and signal receiver clusters are defined and differentially expressed ligands and receptors are identified. To estimate the downstream activation signaling effect of each receptor, we calculate the maximum flow from each receptor to a set of transcription factors using the PPI network. Permutation tests are used to estimate the significance of the receptors and their signaling converging transcription factors. Finally, we calculate an average cluster-to-cluster interaction score and construct a global interaction map per data set.

## Methods

### Data processing, clustering and annotation

The pipeline receives Seurat objects following data processing, normalization, clustering and annotation as was previously described^5^. Briefly, the pipeline consists of the following steps. **LogNormalize**: each feature counts for each cell are divided by the total counts for that cell and multiplied by a scale factor. **Dimensionality reduction:** PCA and tSNE are calculated from the scale normalized data matrix, where each feature normalized expression is scaled across the cells. The number of PCs for the clustering was manually selected based on an elbow plot showing the gain in variance with each additional vector. **Clustering:** First, we calculated the k-nearest neighbors and constructed the KNN graph, in the reduced PCA space. On that graph, a modularity score is optimized using the Leiden clustering method^6^. **Cluster annotation:** was done manually by the use of known cell population markers and projection of known cell-type gene signatures on the tSNE plots.

### Single cell gene signature scoring

Single cell gene signature scoring was used to emphasize the differential expression of ligands and receptors on interacting cell subtypes, as described previously^7^. Briefly, scores were computed by first sorting the normalized scaled gene expression values for each cell followed by summing up the indices (ranks) of the signature genes. A contour plot which takes into account only cells that have a signature score above the indicated threshold was added on top of the tSNE space, to further emphasize the region of high-scoring cells.

### Finding differential expressed receptor-ligands pairs

Ligand and receptor pairs were retrieved from the CellTalkDB database^8^. As a first step, we identified receptors that are differentially expressed in the signal receiving cluster and their corresponding ligands which are differentially expressed in the signal sending cluster, using the Seurat package “FindMarkers” function. We applied the Wilcoxon Sum Rank with limit testing chosen to detect genes that display an average of at least 0.1-fold difference (log-scale) between the two groups of cells and genes that are detected in a minimum fraction of 0.1 in the upregulated group.

### Threshold optimization

To find an optimal threshold, that is balancing between the number of receptors and their activation score, we suggest plotting the mean downstream activation score (DSA) and the number of DE receptors per cluster for each threshold. Thus, the user can identify a range of thresholds that return a significant subset of receptors with a relatively high activation score, for all the clusters in the data.

### Identifying activated transcription factors

To identify potentially activated transcription factors in each cluster, we performed an enrichment test based on the Dorothea TF-gene target database^9^. For a given gene target list, we performed an unpaired Wilcoxon Sum Rank test between the rank distributions of the gene list and the rest of the gene expression vector. Thus we were able to test if the mean expression of the genes in the gene list was taken from the same distribution as the background. The resulting p-values were further FDR corrected.

### Calculation of downstream activation score (DSA)

In addition to the identification of receptors which are differentially expressed, we further calculated their potential downstream activation signaling. Our working assumption is that such a signaling cascade will be reflected in the transcriptomic profile of the cell, and will be likely to converge at a transcription factor downstream.

Firstly, we found transcription factors whose target genes are enriched in each cluster. To avoid the relatively large number of missing gene expression values associated with single cell data (“zero inflated” data) we calculated these parameters per cluster. For each cluster, we considered only those TFs which received an FDR-corrected p-value of 0.05 or lower. Next, we calculated a maximum flow from each receptor towards the enriched layer of transcription factors in a network of protein-protein interactions^8^.

To increase our confidence in the identified signaling pathway, we further normalized the generated network. Each edge received a weight that represented the mutual information between the expression’s distribution in the cluster of genes that made up the edge:

let (*X, Y*) be a pair of random variables, we denote *P_X,Y_*(*x,y*) their joint distribution and *P_X_*, *P_Y_* the margin distribution of *X, Y*. The mutual information of *I*(*X*: *Y*) is defined as:

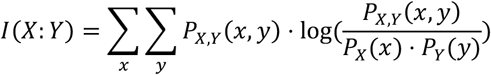

using the mutual information between each pair of genes in the cluster, we marked the edge weight as follows:

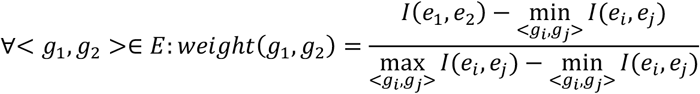

where < *g*_1_, *g*_2_ > is an edge in the PPI network, and *e*_1_, *e*_2_ are the expression vectors in the signal receiver cluster of *g*_1_, *g*_2_ respectively *I*(*e*_1_, *e*_2_) is the mutual information of *e*_1_, *e*_2_. Thus, the weight of a given edge reflects the co-expression of the genes in the cluster, as a proxy for their co-activation.

Following the generation of a weighted signaling network, we added a virtual node to the graph that represents the sink. Each of the identified TFs was then connected to the sink, with an edge weight of infinity. Each edge weight within the resulting normalized network represents its flow capacity. Next, Dinitz’s algorithm^10^ was applied to find the maximum flow from each receptor via the signaling pathway toward the transcription factors ending at the virtual sink node.

To further assess the significance of the flow value, we performed a random degree preserving permutation on the signaling network. For each permutation, we randomly shuffled the edges 10*|E| times, where |E| is the number of edges in the graph, such that each edge will be replaced by an edge with a similar weight. For that, we used the switching algorithm^11^, for each permutation we calculate the max flow, as in the original network, enabling us to provide an empirical statistical value that represents the significance of the observed flow (after FDR correction), compared to flow gained on random networks, while reducing the effect of node centrality in the network.

### Generating a global interactions map

Finally, we calculated the cluster-to-cluster (c1, c2) interaction score for every pair of clusters in the data, using the next formula:

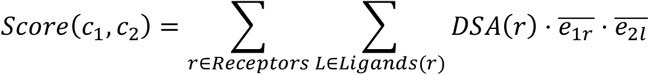

Where *Ligands*(*r*) is the set of ligands in *c*_2_ that corresponds to the receptor *r*, the *DSA*(*r*) is the flow score of receptor *r*, and *e*_1*r*_ is the normalized expression of receptor r in cluster *c*_1_, and *e*_2*l*_ is the normalized expression of ligand *l* in cluster *c*_2_.

## Results

We developed a computational framework, FLOW, that maps the expression of ligand-receptor pairs between different cell populations based on single cell transcriptomes and uses maximum flow computations in a protein-protein interaction network to characterize the downstream pathways from a set of differentially expressed receptors toward dominant TFs.

In order to evaluate our framework we performed systematic validations of each of its steps. Using single cell RNAseq dataset from GL261a murine brain tumor model^4^, we first validated the assumption that upregulated receptors are associated with significant downstream flow, as summarized in **Supp Fig. 1.** Second, we validated the assumption that significant flow values are associated with active transcription factors in each cell population. Third, we generalized this claim and checked if the flow values of all downstream genes were associated with signatures of known activated genes in each cell type. Finally, we evaluated the method’s robustness, by applying it to independent datasets. In the following, we describe these validations and then the application of FLOW to investigate the interaction between Macrophages and CD4+ T cells in the GL261a murine brain tumor model.

### Validation of identified transcription factors

We hypothesized that in most cases, the interaction signal should flow toward a layer of transcription factors, which in turn will propagate the signaling into a cellular response. Therefore, we expected that transcription factors that are up-regulated in a given cluster would also be associated with higher signaling flow values. However, without normalization, the importance value of each TF is strongly associated with the degree of centrality of the node in the PPI network. Thus, we normalized the flow value by the flow value generated in the unweighted PPI network (all capacities are one).

Indeed, we found that there is an association between up-regulated TF and the normalized flow value. Thus, to some extent the TF and its upstream signal transduction pathway are co-regulated. To further validate our result, we tested the correlation between the flow score of a transcription factor and the enrichment of its target genes in the cluster. We assume that enrichment of the TF target genes expression is a strong indicator of TF activation. To this end, we compared the -log (Targets Enrichment P-value) and the normalized flow score of all TFs in each cell type. We found a correlation of 0.3-0.4 for each of the immune cell populations (**Fig. 2B**). Thus, we can further conclude that our flow score contains some indication for the activation of a given transcription factor.

**Figure 2.**
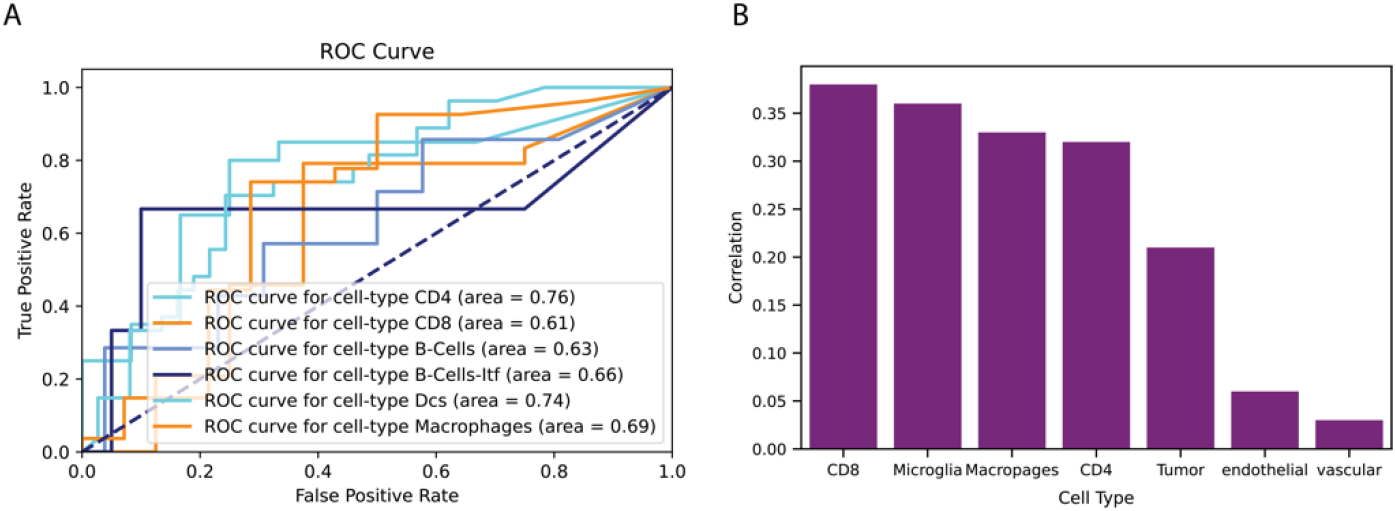
Validation of converging transcription factors. (A) ROC curve analysis of upregulated transcription factor by normalized flow score. (B) Correlation between TFs flow score and the target enrichment score per cell type.

As can be seen in Figure 2, there is a relatively high variance in the quality of performance for different cell populations. This could potentially be explained by the significant bias for commonly studied cell populations, especially in cell-specific TFs and their targets. Thus, commonly studied cell populations are likely to have a relatively larger known subset of TFs and their targets. Therefore, one can expect to observe a better performance of FLOW in those cell populations. Likewise, one can assume that not all cell populations in a biological system are affected in the same manner by cell-cell interactions. Thus, those cells which are not a key interaction axis in the system will yield a lower result by the FLOW analysis.

### Validation of downstream genes scores against MSigDB

To validate our downstream gene scores, we used the MSigDB immune signature database (C7). As each signature in the database is associated with a specific cell type, enrichment analysis for a subset of genes associated with interactions between two specific cell types could be tested. More specifically we performed an enticement analysis of the genes with significant flow values against the entire C7 database. To this end, we first performed a permutation test (as discussed above) for the amount of flow going into each node in the network, nodes/genes with significant flow values were used to perform an enrichment test. Next, we used Fisher Exact Test to examine if the proportion of the gene signatures that were returned from the analysis and are associated with the correct cell types is significantly greater than the overall proportion of the cell type signatures in the database.

Indeed, the proportion of the gene signatures associated with the correct cell types is, in most cases, significantly larger than their proportion in the entire database. Thus, there is an association between genes with significant flow value and biological function in a given cell.

We used the same framework in order to validate our performance against NicheNet^2^. This approach uses Personal Page Rank (PPR) in order to assign an activation score for downstream genes from identified ligands in the data set. The compression demonstrates that the enriched signatures identified using NicheNet were associated with the correct cell type to a lesser degree compared to FLOW (**Supp Fig. 3**).

### Robustness of FLOW

We evaluated the robustness of FLOW and further compared it to the CellChat algorithm^3^. For that, we interactively sampled the data set in fractions of 0.7, 0.5, 0.1, 0.05, and 0.01. For each fraction, we calculated the correlations in interaction scores to the scores gained from the full dataset. We found that FLOW was extremely robust with a drop in correlation observed only with a fraction size of 5%, while CellChat showed a significant drop in correlation at 50% sampling of the data (**Fig. 4**), the entire compression to CellChat is summarized in **Supp Fig. 4.**

**Figure 3.**
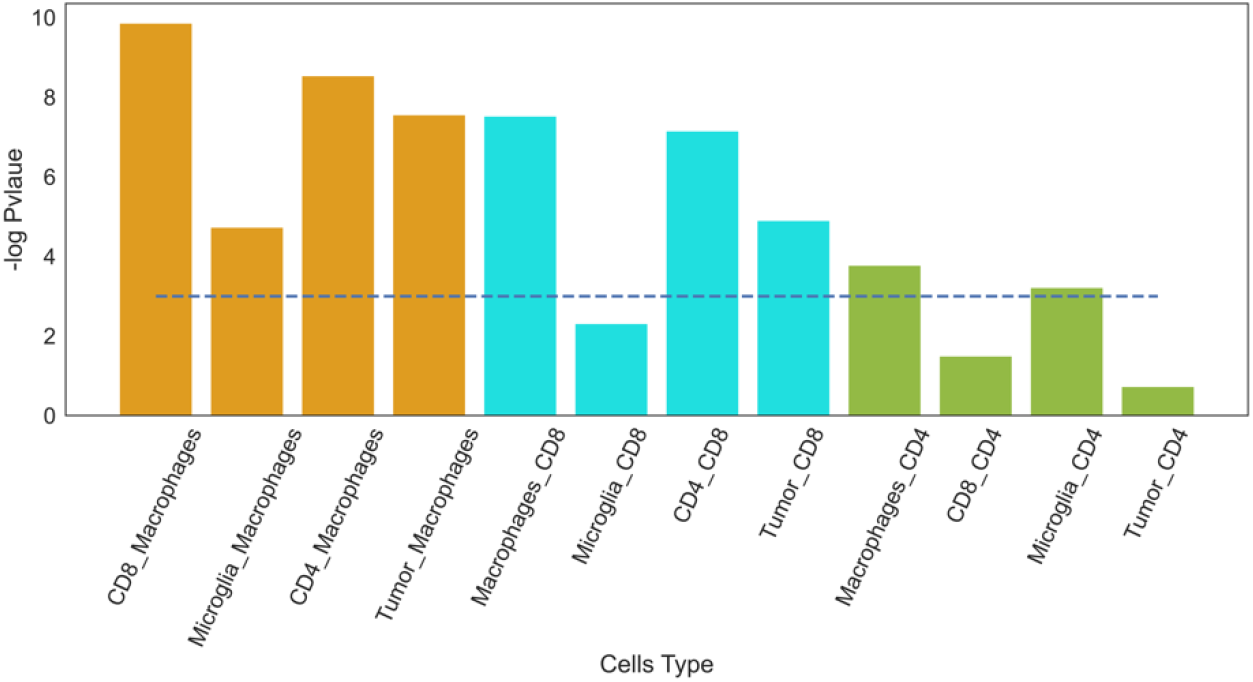
Validation against MSigDB C7 immunological signature database. FLOW was applied to the interaction between each two cell types, and a signature containing significant genes in the receiving cell type was defined. Enrichment of each gene signature associated with the correct corresponding cell type from the MSigDB C7 immunological database was calculated. The blue line represents the significant threshold as -log(0.05)

**Figure 4.**
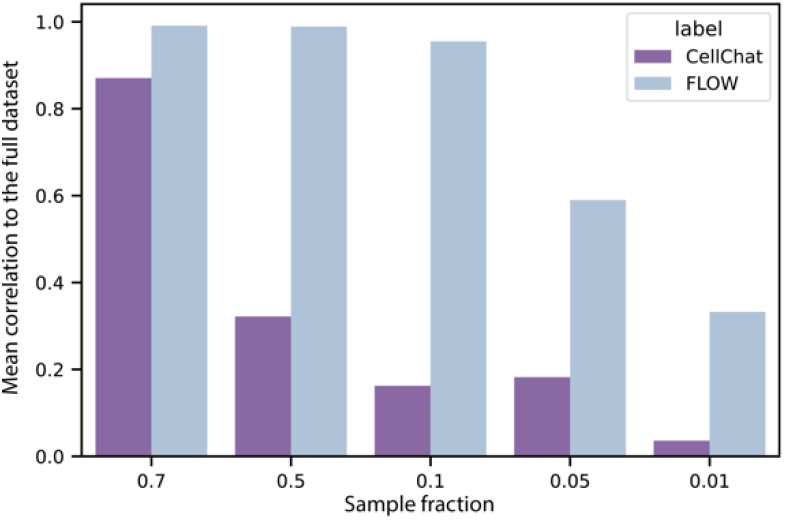
Mean correlation of dataset sub-sampling to the full dataset scores.

We further applied FLOW to a single cell RNA seq data from draining lymph node tissue^33^ and used the same validation framework as discussed above. The results are summarized in **Supp Fig 5.**

### Ligand receptor interactions in the tumor microenvironment

After establishing the accuracy of our method, we set to investigate specific cell-cell interactions inferred from the brain tumor dataset. As was previously shown, increased infiltration of T cells is associated with prolonged survival of GBM patients^12^ and CD4+ T cells play a key role in coordinating antigen-specific immunity through their high plasticity and cytokine-producing ability. However, tumor associated macrophages have been associated with high-grade gliomas and a worsened cancer outcome^31^. It has been proposed that they produce cytokines and other factors to promote a tumor-supportive environment by suppressing the proliferation of anti-tumor CD4+ and CD8+ T cells and promoting the activity of regulatory CD4+ T cells^13^. Interfering with the crosstalk between pro-tumorigenic macrophages and CD4+ T cells has been proposed as a therapeutic strategy, however, detailed information about this interaction is lacking. Thus, out of the multiple cell-cell interactions detected by our method, we used FLOW to investigate the interaction between Macrophages and CD4+ T cells in the GL261a murine brain tumor model.

As a first step, we identified differentially expressed ligand-receptor pairs. While this approach assured us that we are uncovering ligand-receptor pairs that best represent the unique interaction between those clusters, it excluded autocrine interactions or interactions between cells in the same cluster. Analysis of the crosstalk between macrophages and CD4+ T cells clusters identified 37 ligands and 39 corresponding receptors, altogether forming 102 different ligand-receptor interactions (**Fig. 5**). As expected, the signal-receiving cluster is seen to be strongly associated with the receptors while the signal-sender cluster is seen to be associated with the corresponding ligands.

**Figure 5.**
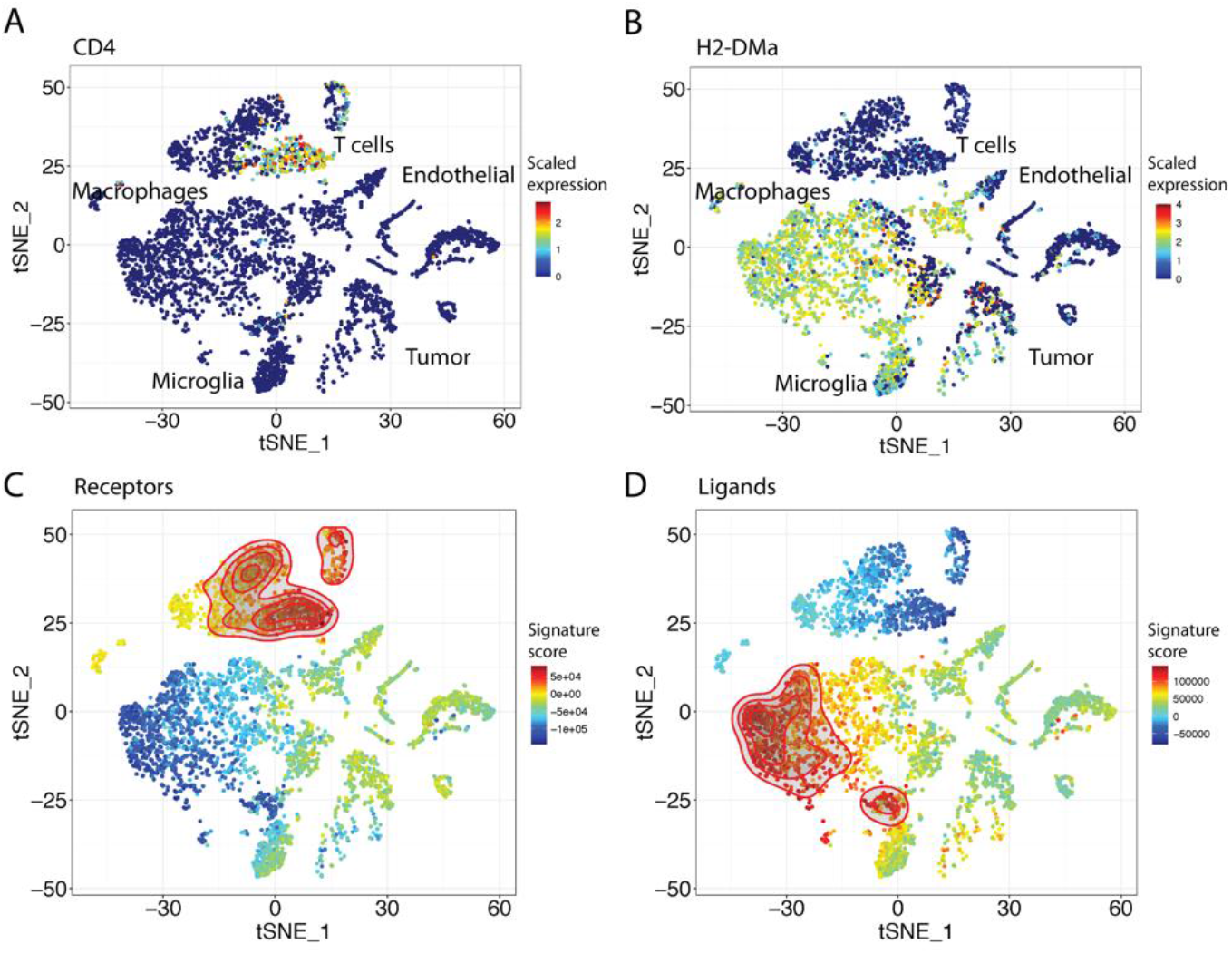
Identification of potential ligand-receptor pairs between macrophages and CD4 T cell clusters. (A-B) The normalized expression projection on the tSNE plot of potential interaction between the MHCII molecule (H2-DMa) on the macrophage cluster and its corresponding receptor CD4 molecule on the T cell cluster. (C-D) Signature projection of 39 receptors identified on the T cell cluster (left) and their 39 corresponding ligands identified on the macrophage cluster (right). The identified receptors are observed to be strongly associated with the T cell cluster while their corresponding ligands are associated with the macrophage and part of the microglia clusters.

Following the identification of ligand-receptor pairs and further calculation of the downstream activation (DSA) score for each of the receptors, the algorithm generates a visualization plot that can be further examined for specific interaction between the two clusters. This plot provides the following information: the many to many ligand-receptor potential interactions, expression level, the significance of differential expression, indication for the downstream signal activation and its statistical significance (**Fig. 6**).

**Figure 6.**
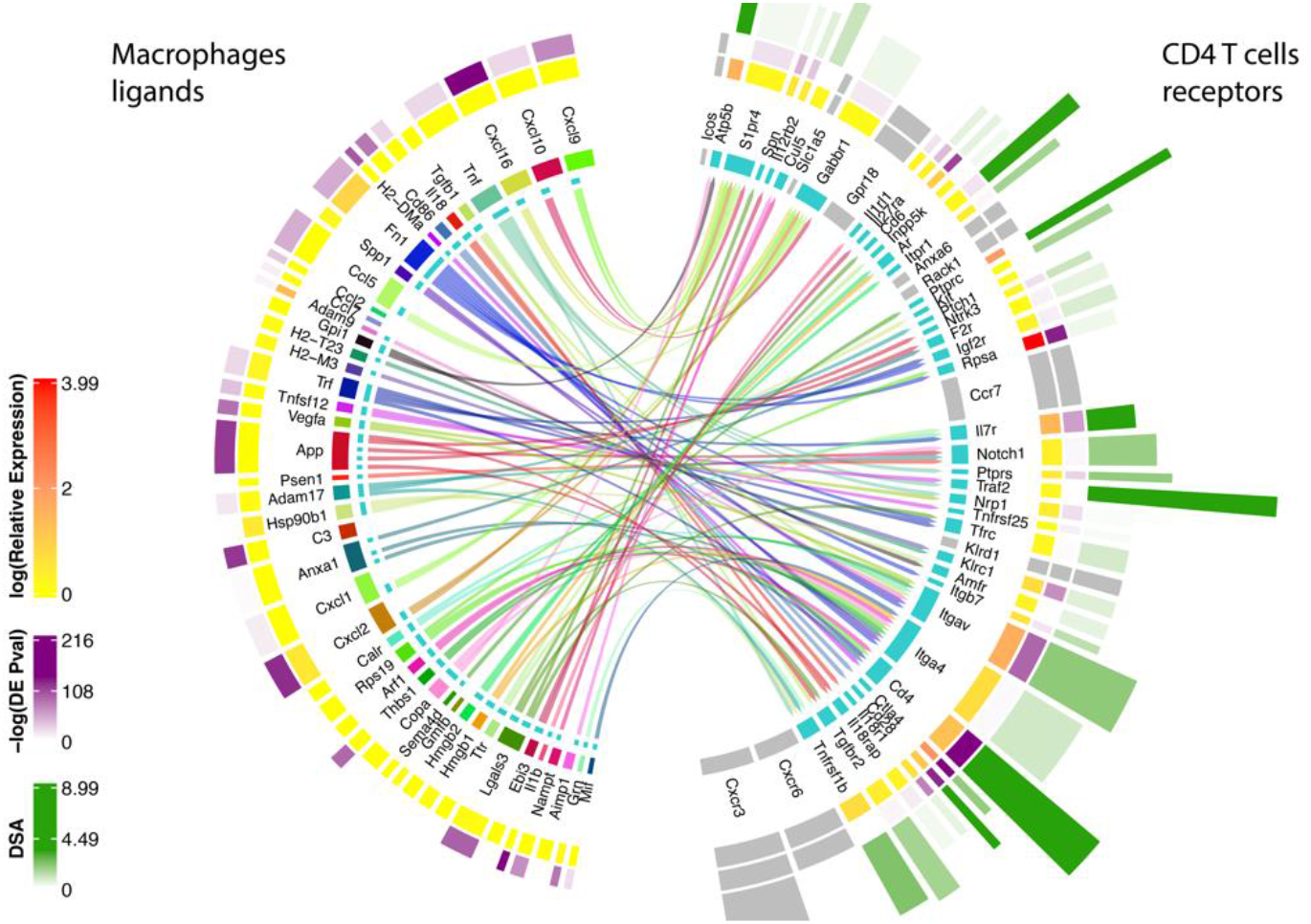
Ligand-receptor ‘ interaction with integrated receptor downstream activation signaling score. Detailed analysis of the differentially enriched ligand-receptor interactions between macrophage (signal-sender) and CD4+ T cell (signal-receiving) clusters. Inner lines indicate potential ligand-receptor connections and the width of the inner circle ribbon indicates the number of potential connections. The second circle ribbon reflects the expression level, the third outer ribbon indicates the p-value of upregulation compared to all other clusters in the dataset and the outer ribbon indicates the downstream activation score (DSA). Receptors that did not show significant value in the permutation test are colored in grey.

Out of these multiple significant ligand-receptor interactions, here we focus on two opposite potential interactions between macrophages and CD4 T cells, which further highlights the complexity of signals a cell receive in this microenvironment. On the one hand, a pro-inflammatory interaction through the IL18 receptor and its ligand IL18a, leads to type I activation in the form of IFNγ synthesis from Th1 cells^14^. On the other hand, an immunosuppressive interaction through the TGF-β receptor and its ligand TGF-β. Moreover, signaling via TGF-β can subvert T-cell immunity by favoring regulatory T-cell differentiation, further reinforcing immunosuppression within tumor microenvironments^15^. Another example is the two interactions between co-stimulatory CD28 and co-inhibitory CTLA4 which compete on the same ligand CD86.

### Signaling converging transcription factors in CD4 T-cells

As part of the framework, FLOW highlights significant transcription factors that are likely to be activated by the signals received from multiple receptors. To this end, we applied the multi-source max flow algorithm, from all the receptors down to the defined set of transcription factors and calculated the amount of flow that is going through each of the transcription factors. Next, to rank the contribution of the different TFs, we calculated the importance coefficient of the TF (**Fig. 7A**). Specifically, we removed each of the TFs from the network, and recalculated the multi-sourced max flow in order to obtain the difference in the maximum flow in the network. This yielded a subnetwork that represents the flow pathways from multiple receptors toward a specific transcription factor, allowing us to further investigate a specific signal transduction pathway and detect potential hubs (**Fig. 7B**).

**Figure 7.**
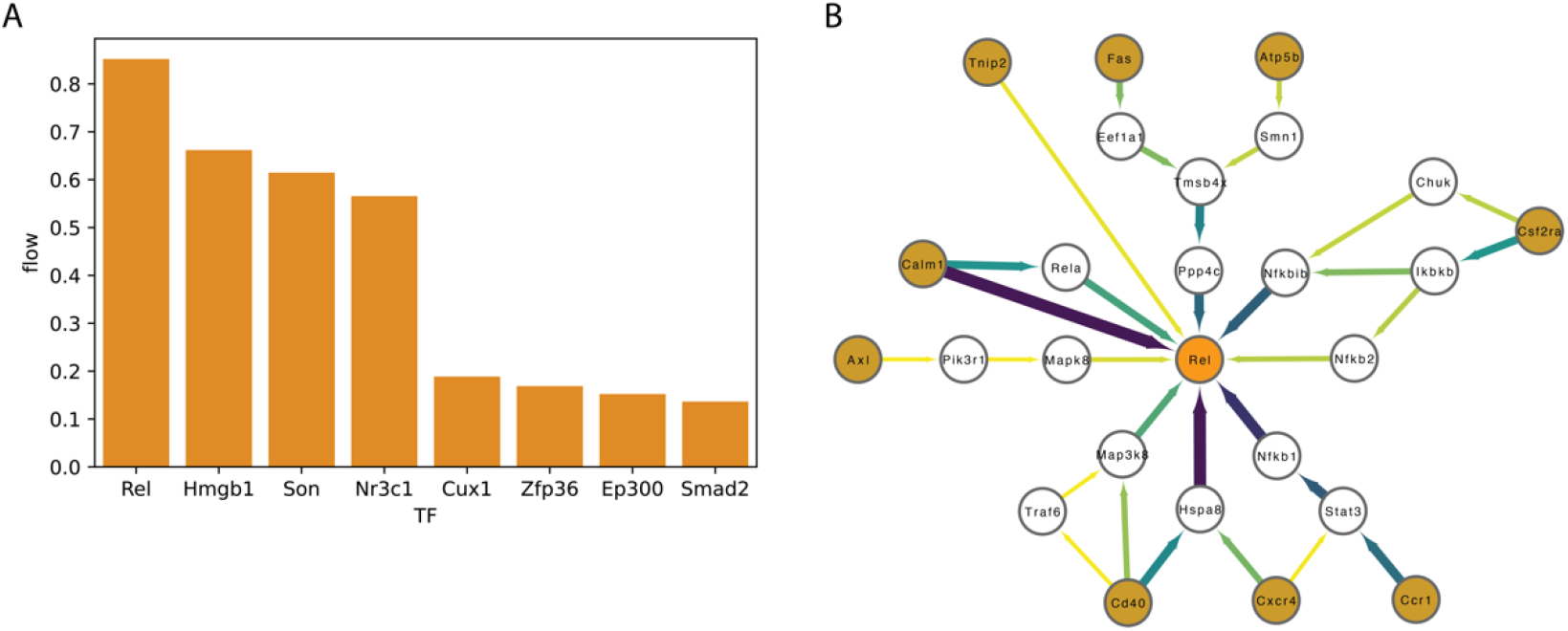
Identification of downstream signaling converging transcription factors for CD4 T cells. (A) A bar plot showing the top-ranking TFs in the interaction between macrophage and CD4 T cell clusters shown in Fig. 6. (B) Subnetwork of the flow from multiple receptors (orange) to the Proto-Oncogene, NF-KB Subunit (Rel) transcription factor. The edge color and width represent the amount of flow that is passing through the edge as a proxy for the significance of the pathway in the subnetwork.

The top-ranking transcription factor, c-Rel, plays a key role in mediating the proliferation, differentiation, and cytokine production of T cells. This protein is crucial for optimal IL-2 production and expression of IL-2Rα (CD25) in T cells^16^. The second top-ranking Hmgb1 has also been shown to be a key proliferative signal for CD8 and CD4 T cells, as well as for the activation of CD4 cells. While the importance of the identified TFs to the survival, differentiation and cytotoxic capacity has been previously demonstrated, here we show that these TFs play a role in the specific interaction between macrophages and CD4 tumor infiltrating lymphocytes (TILs) in the context of the brain tumor microenvironment^17^.

### Pathways enrichment analysis following macrophages and CD4 T-cells interaction

Next, we used FLOW to detect enriched pathways between those two cell populations. We performed an enrichment test, as discussed above against KEGG and GO databases. In order to eliminate biases towards a specific cell population, expressed genes in each cluster were used as background for the enrichment test. Indeed, testing for enrichment with a random set of genes did not yield any significant results.

In the KEGG analysis, we observed an enrichment of immune and T cells related pathways, such as “Th1 and Th2 cell differentiation”, “Pathways in cancer” and “T cell receptor signaling pathway” as well as, pathways that are associated with a specific conditions such as “Human T-cell leukemia virus 1 infection”.

In the GO analysis, the most significantly enriched terms found were strongly associated with T-cells activation and immune response, such as “lymphocytes activation”, “response to cytokines” and “immune response”, indicating that genes with a significant amount of flow, are associated with the expected function of the cells.

The entire result of the KEGG and GO enrichment analysis can be found in **Supp Tables 1 & 2.**

### Global interactions in the tumor microenvironment

By applying the previously described method on each pair of clusters in the data set, we have generated a global map of interactions between all cell populations in the dataset (**Fig. 8A**). We hypothesized that generating such an interaction map could help us to better understand the global dynamics between different cell populations in the data set, and uncover the key interaction axes common and cell-type specific to a studied biological system (**Fig. 8B**).

**Figure 8.**
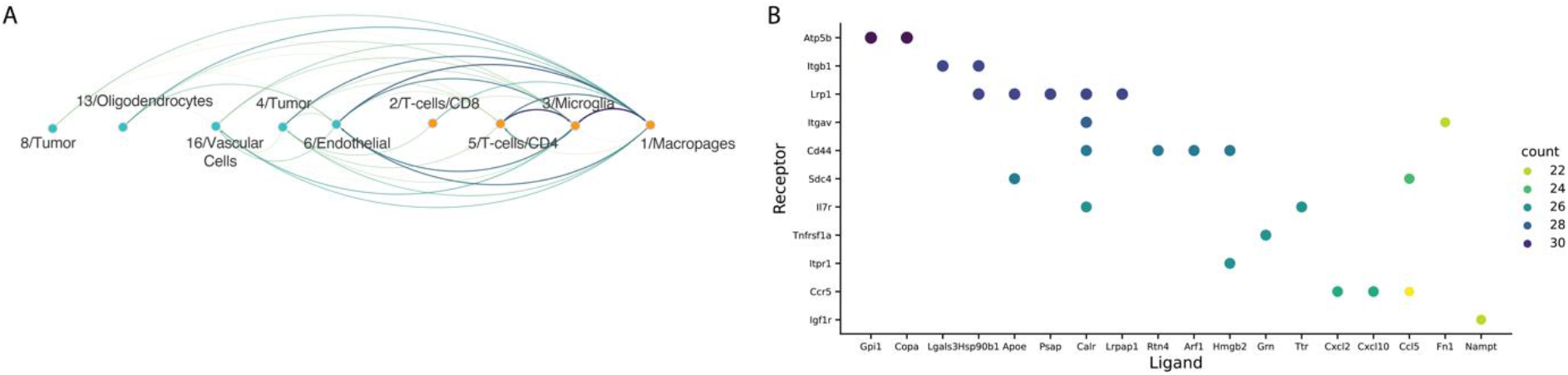
Global interaction map. (A) Global interaction plot, demonstrating interactions between the different clusters in the dataset. Edges radiate from the signal sender cluster to the signal receiving cluster, edge colors and width represent the strength of the interaction. (B) Top 25 ligand-receptor interactions that were identified as active between multiple cell types in the dataset.

As can be observed in Figure 8B, there are almost no significant interactions between non-immune cell populations (endothelial, vascular, tumor and oligodendrocyte). However, they do show relatively strong interactions with antigen-presenting myeloid cells (macrophages and microglia). In addition, we detected expected interactions between CD4+ helper T cells and the antigen-presenting populations.

Finally, our global analysis can also be used to highlight common ligand-receptor interactions between different cell types. Indeed, we found many such interactions which are common to communications between multiple cell types.

Taken together, our global analysis was well adapted to capture some of the known cell-cell interactions in this microenvironment and highlighted unique and more common interactions that must be taken into account in perturbation applications.

## Discussion

The task of quantitative modeling of cell-to-cell interaction based on transcriptomic data and single cell transcriptomics specifically is heuristic by nature. First, gene expression data is not able to detect direct interaction between different cells, which leaves us only with the detection of the interaction potential. Second, gene expression also does not necessarily reflect gene activation. Third, the relatively low mRNA capture rate of the single cell RNAseq technologies generates “zero-inflated” datasets, which makes it harder to detect such complex events per cell.

In this article, we proposed a computational framework view of the maximum flow problem. Assuming that a significant proportion of the signals received by the cell via its receptors will propagate downstream toward a transcription factor. Provided the right cues, this transcription factor will in turn affect a vast number of genes. Thus, an analog to maximum flow in a network enables us not only to uncover important ligand-receptor pairs participating in cell-cell crosstalk but also, to detect likely pathways by which the signaling occurs and the transcription factors that aggregate the signals from different receptors. Importantly, by providing clear start and end points for a signaling pathway, the suggested framework avoids making prior assumptions on the pathway length or limiting the analysis to a single pathway. This approach enables us to model complex interaction systems, such as the immune response in the tumor microenvironment, across a relatively large subset of channels in each cell type.

To further demonstrate the power of FLOW, we applied it to a single cell RNA seq obtained from an independent Glioma brain tumor mouse model containing both wild-type and myeloid MHCII knockout mice^32^. Using FLOW, we identified multiple potential interactions between various cell types some of which were differential between the WT and the KO cells. Spp1-Itgb1 was found to be a key communication axis between macrophages and CD8+ T cells in the KO samples. Furthermore, our method identified that the downstream signaling pathways in the interaction between these two cell types converged at the transcription regulator Nfat2, which is a positive regulator of Tox (critical regulator of T cell exhaustion). Complementary biological experiments were able to validate the effect of Spp1-Nfat2-Tox axis as was predicted by our method.

Cell-cell interaction involves the activation of multiple receptors followed by complex intracellular signal transduction. These signals must be aggregated for the cell to reach a reaction decision, thus highlighting the role of key transcription factors as a layer that aggregates the different signals into a cellular response is unique to our approach. Further characterization of transcription factors identified in a specific biological setting is, however, needed. Nevertheless, this approach opens new therapeutic avenues for the perturbation of a cellular response following cell-cell interactions.

Similar to our framework, CellCall^18^ assumes that signaling via receptor leads to transcription factor activation. The CellCall model assigns an activation score for a set of KEGG pathways while also taking into account the enrichment of TF’s target genes in each pathway. Rather than looking at a set of disjoint pathways, our suggested FLOW approach takes advantage of an interaction network. This allows the discovery of novel activation pathways related to specific cell populations and biological conditions. Furthermore, our maximum flow framework can model the underline assumption that signals from different receptors are converging into a subset of dominant transcription factors. Lastly, our methodology could assign activation scores for transcription factors based on network importance rather than a single pathway. In this article, we have shown that the amount of flow of a given receptor is associated with its upregulation in different cell populations. The signal flow was also found to be associated with the upregulation of the transcription factor predicted to further converge the signals. Thus, our interaction potential between two clusters includes not only the expression of the receptor and ligand but its entire signal transduction pathway.

On a global scale, our framework allows us to inspect the interaction map of an entire single cell data set. Indeed, various microenvironments are characterized by multiple cell-type populations that constantly interact with each other. Such complex and dynamic interaction as well as a comparison between different states of the system could greatly benefit from such global analysis.

It is important to note, that the activation of a signaling pathway in biological systems usually also includes post-translational modifications, which are not measured by any form of RNA seq data. Thus, any activation score that is based solely on RNA expression will never capture the entire pathway activation process.

Using data sets taken from a tumor microenvironment, we have shown that our method can reflect dynamics in the data that agreed with known biological assumptions. To reduce the effect of the data sparsity associated with single cell RNAseq most of our calculations are done at the cluster level. However, we believe that our suggested framework, with the proper normalization, may enable us to assign an interaction potential score to every single cell in the dataset, and by that reveal more complex biological dynamics inside the clusters. Combining such cell-specific scores with other single cell analysis approaches will allow us to ask more complex questions, such as how the interactions can affect the cell differentiation trajectories, and understand how cell-to-cell interactions change in different biological conditions.

## Supplementary

**Supplementary Table 1.** KEGG Pathway enrichment analysis Top 40 pathways.

| Gene_set Term | Overlap | P-value | Adjusted P-value | Old P-value | Old Adjusted P-value | Odds Ratio | Combined Score |
| --- | --- | --- | --- | --- | --- | --- | --- |
| KEGG_202 Th1 and Th2 cell differentiation | 29/92 | 3.91E-32 | 8.14E-30 | 0 | 0 | 37.72301587 | 2728.141125 |
| KEGG_202 Pathways in cancer | 54/531 | 7.11E-32 | 8.14E-30 | 0 | 0 | 10.13813076 | 727.1197882 |
| KEGG_202 Viral carcinogenesis | 37/203 | 3.51E-31 | 2.68E-29 | 0 | 0 | 18.79686851 | 1318.122894 |
| KEGG_202 Th17 cell differentiation | 27/107 | 4.83E-27 | 2.77E-25 | 0 | 0 | 27.40583678 | 1660.638924 |
| KEGG_202 Human T-cell leukemia virus 1 infection | 34/219 | 2.79E-26 | 1.28E-24 | 0 | 0 | 15.28611846 | 899.4451359 |
| KEGG_202 Hepatitis B | 30/162 | 1.23E-25 | 4.69E-24 | 0 | 0 | 18.63731457 | 1069.024579 |
| KEGG_202 T cell receptor signaling pathway | 24/104 | 3.88E-23 | 1.27E-21 | 0 | 0 | 24.06244898 | 1241.700294 |
| KEGG_202 PD-L1 expression and PD-1 checkpoint pathway in cancer | 22/89 | 5.15E-22 | 1.47E-20 | 0 | 0 | 26.14103571 | 1281.388054 |
| KEGG_202 Epstein-Barr virus infection | 28/202 | 2.02E-20 | 5.14E-19 | 0 | 0 | 13.05852053 | 592.1960032 |
| KEGG_202 Kaposi sarcoma-associated herpesvirus infection | 27/193 | 7.79E-20 | 1.78E-18 | 0 | 0 | 13.14983073 | 578.585421 |
| KEGG_202 Yersinia infection | 23/137 | 6.90E-19 | 1.44E-17 | 0 | 0 | 16.08868207 | 672.787555 |
| KEGG_202 Osteoclast differentiation | 22/127 | 2.01E-18 | 3.73E-17 | 0 | 0 | 16.64823597 | 678.3476072 |
| KEGG_202 Human papillomavirus infection | 32/331 | 2.12E-18 | 3.73E-17 | 0 | 0 | 8.775016581 | 357.1114678 |
| KEGG_202 Human immunodeficiency virus 1 infection | 26/212 | 1.10E-17 | 1.80E-16 | 0 | 0 | 11.2431966 | 439.0255868 |
| KEGG_202 TNF signaling pathway | 20/112 | 3.99E-17 | 6.09E-16 | 0 | 0 | 17.14597521 | 647.4372133 |
| KEGG_202 Human cytomegalovirus infection | 26/225 | 4.88E-17 | 6.99E-16 | 0 | 0 | 10.50172674 | 394.4307175 |
| KEGG_202 Measles | 21/139 | 2.25E-16 | 3.03E-15 | 0 | 0 | 14.07439174 | 507.1186762 |
| KEGG_202 JAK-STAT signaling pathway | 22/162 | 4.35E-16 | 5.54E-15 | 0 | 0 | 12.46390977 | 440.8566196 |
| KEGG_202 FoxO signaling pathway | 20/131 | 9.62E-16 | 1.16E-14 | 0 | 0 | 14.19732986 | 490.9051936 |
| KEGG_202 PI3K-Akt signaling pathway | 29/354 | 7.34E-15 | 8.40E-14 | 0 | 0 | 7.215051282 | 234.8178434 |
| KEGG_202 Natural killer cell mediated cytotoxicity | 19/131 | 1.35E-14 | 1.48E-13 | 0 | 0 | 13.31289286 | 425.1151017 |
| KEGG_202 AGE-RAGE signaling pathway in diabetic complications | 17/100 | 2.28E-14 | 2.37E-13 | 0 | 0 | 15.96940142 | 501.6432303 |
| KEGG_202 Adherens junction | 15/71 | 2.69E-14 | 2.68E-13 | 0 | 0 | 20.74838301 | 648.3035846 |
| KEGG_202 C-type lectin receptor signaling pathway | 17/104 | 4.49E-14 | 4.28E-13 | 0 | 0 | 15.23207444 | 468.1612383 |
| KEGG_202 Inflammatory bowel disease | 14/65 | 1.49E-13 | 1.37E-12 | 0 | 0 | 21.18569781 | 625.6539522 |
| KEGG_202 Lipid and atherosclerosis | 22/215 | 1.71E-13 | 1.51E-12 | 0 | 0 | 9.01671876 | 265.0472994 |
| KEGG_202 MAPK signaling pathway | 25/294 | 2.52E-13 | 2.14E-12 | 0 | 0 | 7.412852703 | 215.0494666 |
| KEGG_202 Focal adhesion | 21/201 | 4.12E-13 | 3.37E-12 | 0 | 0 | 9.197379032 | 262.2917197 |
| KEGG_202 Proteoglycans in cancer | 21/205 | 6.07E-13 | 4.79E-12 | 0 | 0 | 8.995595196 | 253.0484462 |
| KEGG_202 Apoptosis | 18/142 | 7.39E-13 | 5.64E-12 | 0 | 0 | 11.33935227 | 316.7427708 |
| KEGG_202 Wnt signaling pathway | 19/166 | 1.08E-12 | 8.00E-12 | 0 | 0 | 10.12506122 | 278.9648235 |
| KEGG_202 Leishmaniasis | 14/77 | 1.78E-12 | 1.28E-11 | 0 | 0 | 17.13986928 | 463.6829503 |
| KEGG_202 Shigellosis | 22/246 | 2.65E-12 | 1.84E-11 | 0 | 0 | 7.756542799 | 206.7724463 |
| KEGG_202 NF-kappa B signaling pathway | 15/104 | 9.46E-12 | 6.19E-11 | 0 | 0 | 13.0332655 | 330.8425913 |
| KEGG_202 Toll-like receptor signaling pathway | 15/104 | 9.46E-12 | 6.19E-11 | 0 | 0 | 13.0332655 | 330.8425913 |
| KEGG_202 Pertussis | 13/76 | 2.53E-11 | 1.61E-10 | 0 | 0 | 15.85342262 | 386.8038732 |
| KEGG_202 Cellular senescence | 17/156 | 3.82E-11 | 2.37E-10 | 0 | 0 | 9.50850748 | 228.0809808 |
| KEGG_202 Hepatitis C | 17/157 | 4.24E-11 | 2.55E-10 | 0 | 0 | 9.44010771 | 225.4715921 |
| KEGG_202 Growth hormone synthesis, secretion and action | 15/119 | 6.85E-11 | 4.02E-10 | 0 | 0 | 11.14495003 | 260.8321491 |
| KEGG_202 Chemokine signaling pathway | 18/192 | 1.25E-10 | 6.99E-10 | 0 | 0 | 8.060310482 | 183.7752513 |
| KEGG_202 Transcriptional misregulation in cancer | 18/192 | 1.25E-10 | 6.99E-10 | 0 | 0 | 8.060310482 | 183.7752513 |

**Supplementary Table 2.**
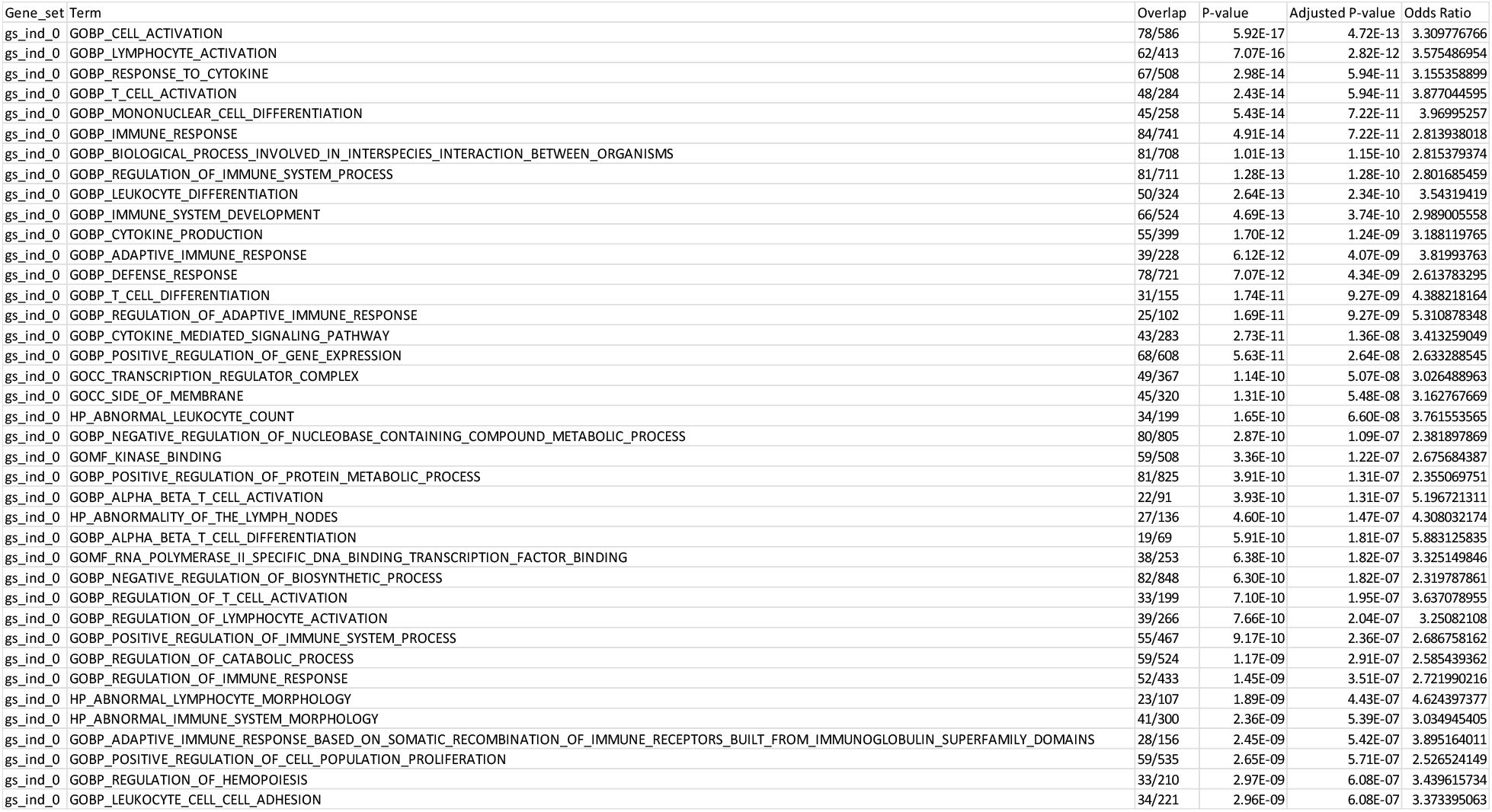
GO Terms enrichment analysis Top 40 Terms.

### Validation of up-regulated receptor

We checked if receptors that are up-regulated in a given cluster are associated with high flow values defined by our generated network. Thus, not only is the receptor itself upregulated but also its downstream signaling pathway.

To this end, for each cluster, we identified a set of receptors that are upregulated relative to all other clusters in the data set. We then defined an equally balanced list of receptors that are upregulated in the cluster and receptors which are not upregulated. For each receptor, we calculated the max flow value, and used these values to calculate ROC AUC of the classification of upregulated receptors (**Supp Fig.1**).

**Supplementary Figure 1.**
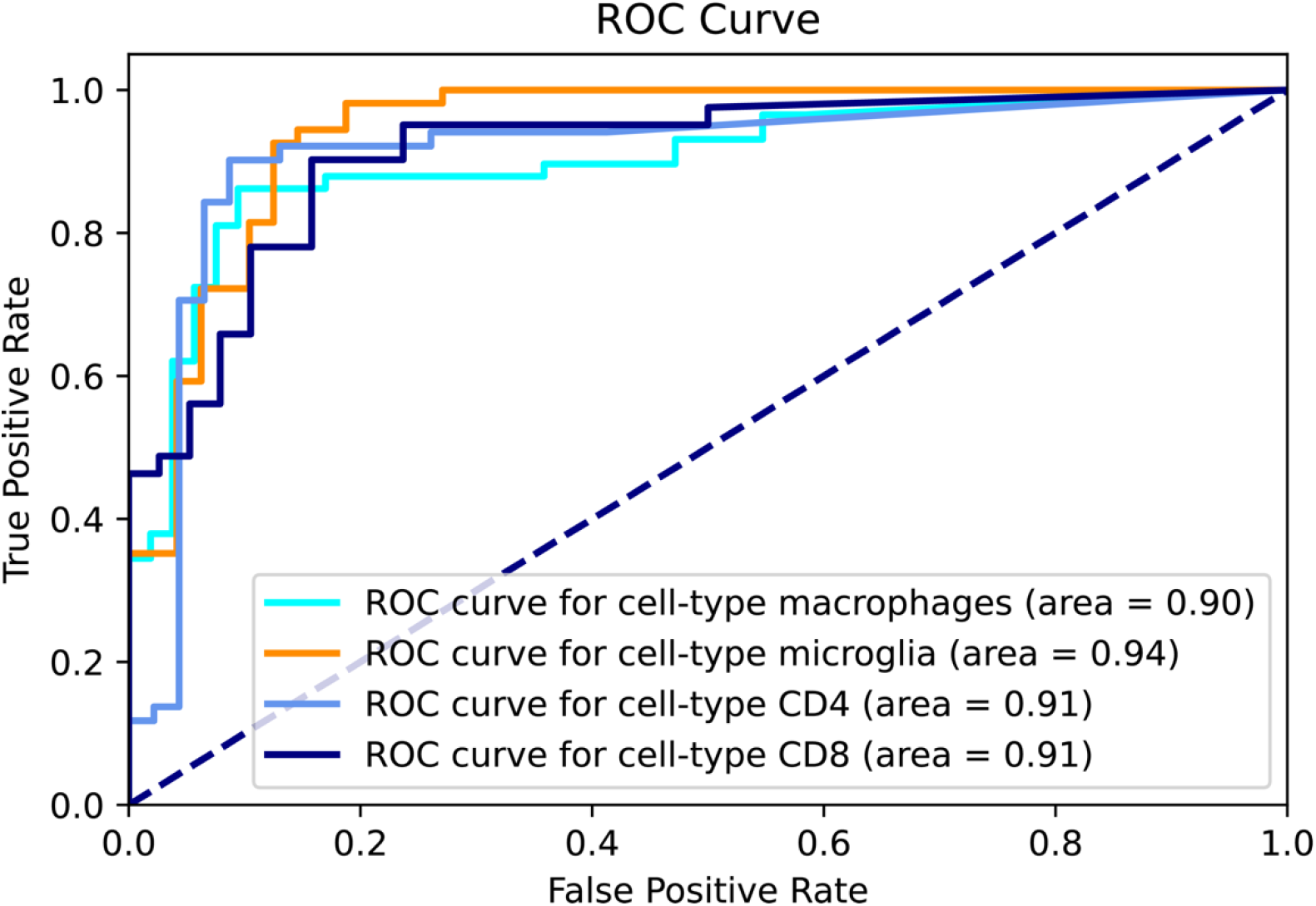
Validation of receptors detection. ROC curve analysis of upregulated receptors by flow shows that flow values are significantly associated with upregulated receptors.

### Global interaction quantification

In the global analysis of the current dataset, we expect to find stronger interactions to or from immune compared to non-immune clusters. To quantify such a trend, we calculated the mean interaction score of immune-related interactions (at least one of the clusters in the interaction represents immune cell type), to the mean score of non-immune interaction (**Supp Fig 2.**). As expected, the immune interaction score was indeed stronger.

**Supplementary Figure 2.**
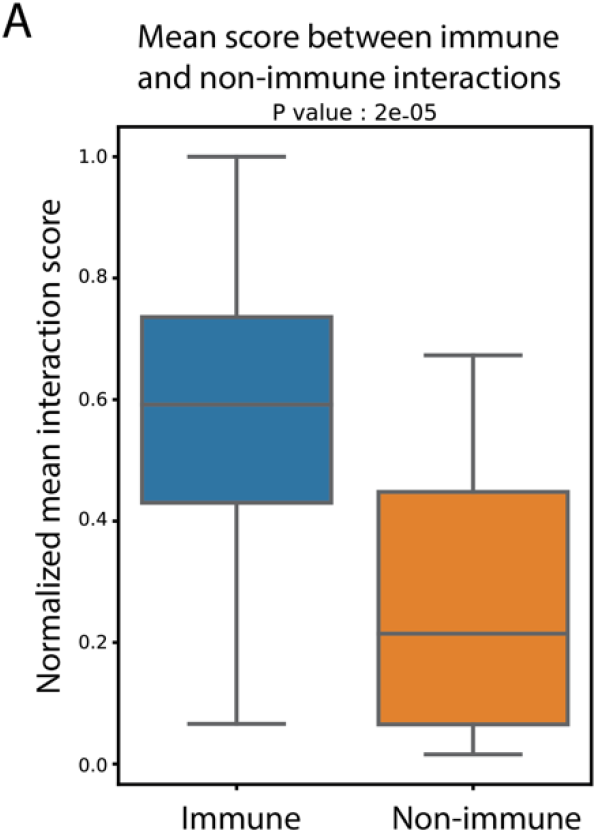
Validation of global immune and non-immune interactions and method robustness. (A) Normalized mean interaction score of immune cell types compared to interactions between non-immune cell types, p-value was calculated using t-test statistic.

### NicheNet MSigdb compression

We run our MSigdb validation for the downstream signal return from NicheNet. First, the activation score of each gene was calculated for each ligand returned by the method. Next, we summed the activation score of each gene over all the identified ligands, then we choose the top n genes (where n is the number of genes returned by FLOW) with the highest score to the enrichment analysis, the results are presented in **Supp Fig 3.**

**Supplementary Figure 3.**
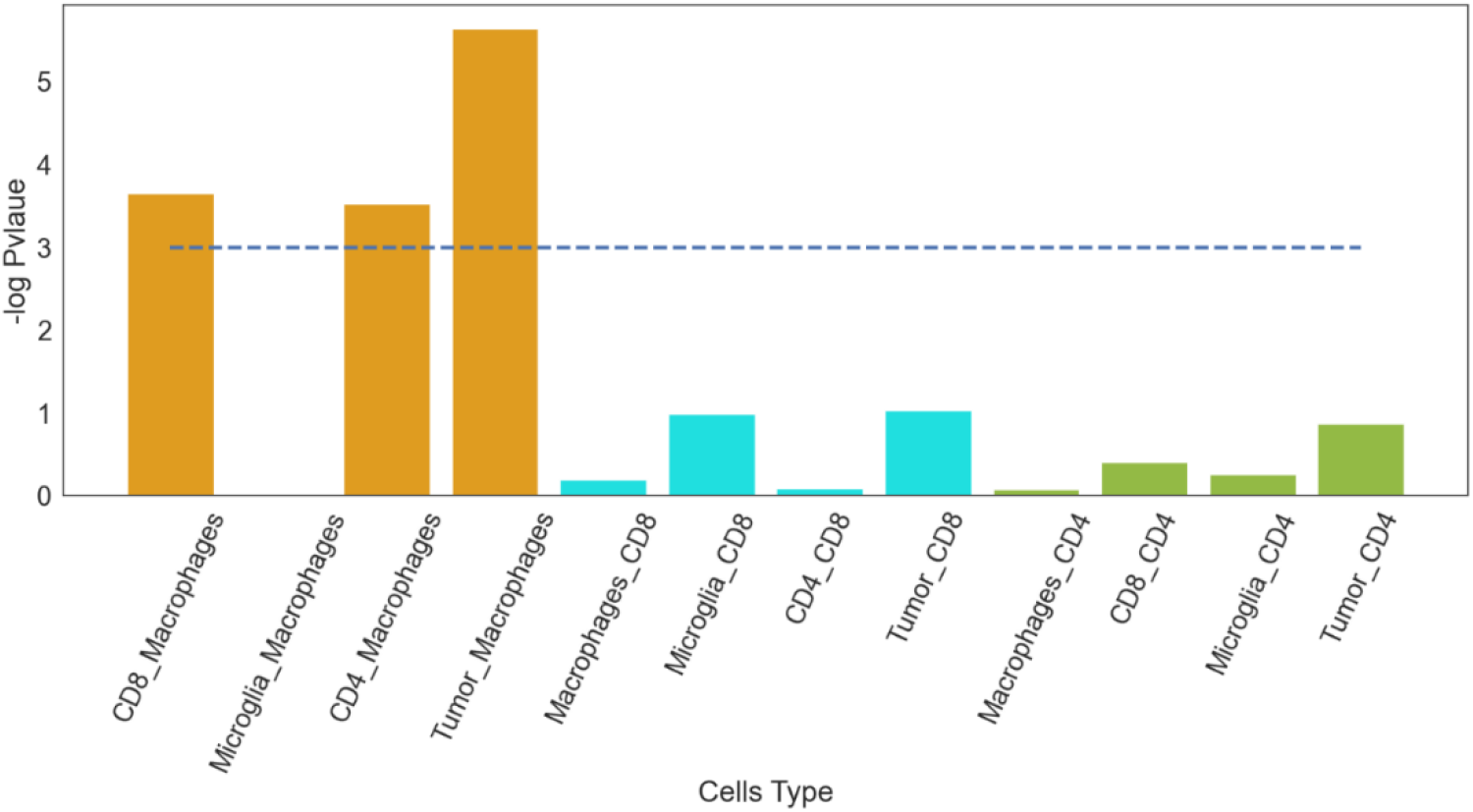
Validation of NicheNet against MSigDB C7 immunological signature database.

### Comparison to CellChat method

We compared our results to the recently published CellChat algorithm. As discussed previously, comparing the identification processes of cluster-specific receptor-ligand pairs using different tools is a non-trivial task. CellChat provides cell-cell communication on a global scale, whereas we were aiming to see if the same biological trends can be observed using the CellChat toolkit, as we observed by FLOW. We generated all-to-all interactions using CellChat, and compared the Immune interactions to the non-immune interactions score. CellChat mean score of the immune interaction was significantly higher than the non-immune interaction (p-value = 0.03), which is thus in agreement with the FLOW method (**Supp Fig. 4**).

As we cannot compare the methods to a “gold standard” data set, we next compared the relationship between the methods’ scores. To this end, we compared FLOW to the score of each receptor. We discovered a significant correlation of 0.4 between our flow scores and CellChat communication probability.

Furthermore, we compared our flow permutation score between receptors that were common to both methods and receptors that were not. We found a significant difference in the mean score value between the two groups (p-value = 0.006), and ROC AUC of 0.7.

**Supplementary Figure 4.**
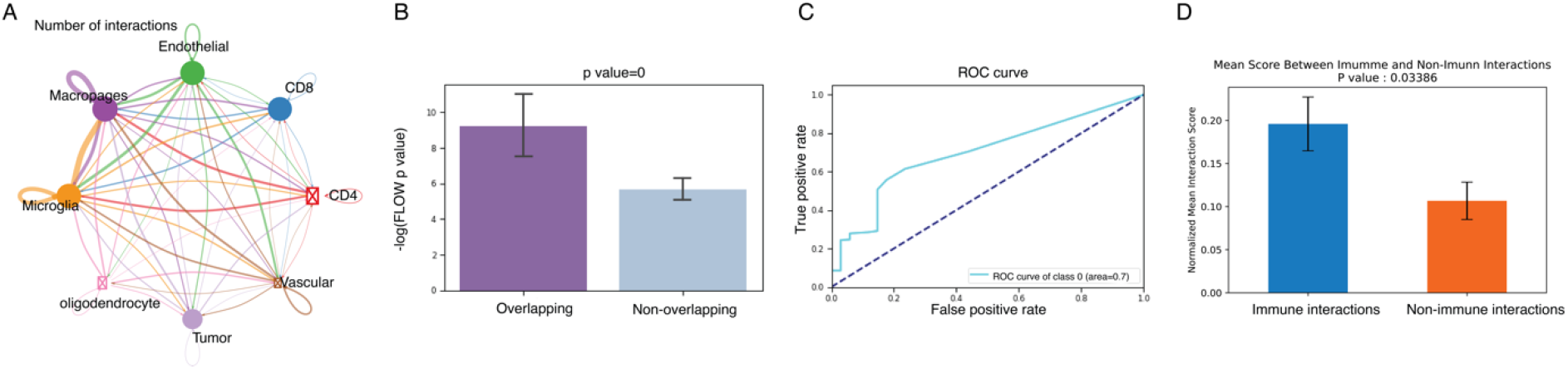
Comparison to the CellChat method. (A) CellChat global interaction map. (B) Mean score of overlapping and non-overlapping interactions between the methods. (C) ROC AUC results for the overlapping interactions by the FLOW score. (D) Normalized mean interactions score of immune cell types compared to interactions between non immune cell types, p value was calculated using t test statistic.

### Lymph node validation

To check the robustness of our framework, we ran the flow pipeline on the published Single Cell RNA seq dataset of a lymph node), a summary of the results is shown in **Supp Fig** 5.

**Supplementary Figure 5.**
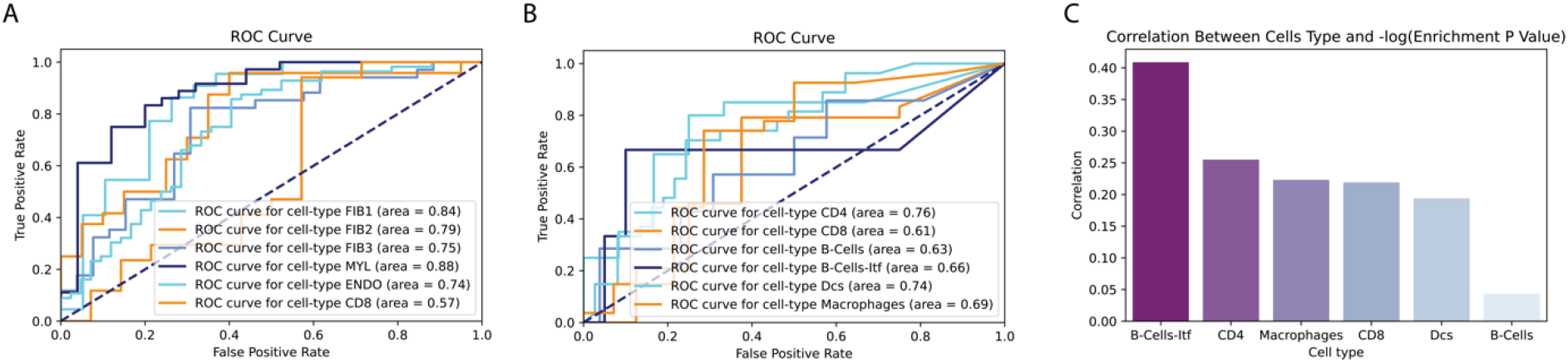
Validation on Lymph node dataset. (A) ROC curve analysis of upregulated receptors by FLOW. (B) ROC curve analysis of upregulated transcription factors by normalized FLOW score. (C) Correlation between TFs FLOW score and the target enrichment score per cell type.

